# HLA-class II specificity assessed by high-density peptide microarray interactions

**DOI:** 10.1101/2020.02.28.969667

**Authors:** Thomas Osterbye, Morten Nielsen, Nadine L. Dudek, Sri H. Ramarathinam, Anthony W. Purcell, Claus Schafer-Nielsen, Soren Buus

**Affiliations:** Department of Immunology and Microbiology, University of Copenhagen, Denmark; Department of Health Technology, Technical University of Denmark, Lyngby, Denmark; Department of Biochemistry & Molecular Biology and Infection and Immunity Program, Biomedicine Discovery Institute, Monash University, Clayton VIC 3800, Australia; Schafer-N, Copenhagen, Denmark; Instituto de Investigaciones Biotecnológicas, Universidad Nacional de San Martín, San Martín, Argentina

## Abstract

The ability to predict and/or identify MHC binding peptides is an essential component of T cell epitope discovery; something that ultimately should benefit the development of vaccines and immunotherapies. In particular, MHC class I (MHC-I) prediction tools have matured to a point where accurate selection of optimal peptide epitopes is possible for virtually all MHC-I allotypes; in comparison, current MHC class II (MHC-II) predictors are less mature. Since MHC-II restricted CD4+ T cells control and orchestrate most immune responses, this shortcoming severely hampers the development of effective immunotherapies. The ability to generate large panels of peptides and subsequently large bodies of peptide-MHC-II interaction data is key to the solution of this problem; a solution that also will support the improvement of bioinformatics predictors, which critically relies on the availability of large amounts of accurate, diverse and representative data. Here, we have used recombinant HLA-DRB1*01:01 and HLA-DRB1*03:01 molecules to interrogate high-density peptide arrays, *in casu* containing 70,000 random peptides in triplicates. We demonstrate that the binding data acquired contains systematic and interpretable information reflecting the specificity of the HLA-DR molecules investigated. Collectively, with a cost per peptide reduced to a few cents combined with the flexibility of recombinant HLA technology, this poses an attractive strategy to generate vast bodies of MHC-II binding data at an unprecedented speed and for the benefit of generating peptide-MHC-II binding data as well as improving MHC-II prediction tools.

## Introduction

The binding of peptides to major histocompatibility complex class II (MHC-II) molecules is one of the most selective events in antigen presentation. Expressed by professional antigen presenting cells and presenting exogenous peptides to CD4+ T cells, MHC-II is central to adaptive immunity. Since CD4+ T cells, restricted to MHC-II, are recognized for their orchestrating role in shaping both humoral and cellular immune responses, much effort has been put into understanding the specificity of MHC-II. The human MHC (human leucocyte antigen) HLA loci is one of the most polymorphic gene families known (1), and as of September 2019, the number of HLA class II HLA-II alleles amounts to 7,065 according to hla.alleles.org, not considering the additional diversity created by the combinatorial HLA-DPA/DPB and HLA-DQA/DQB pairing. Further, considering the number of peptides that can be presented and the huge diversity of T cell receptors that can recognize peptide-HLA-II (pHLAII) complexes, the combinations of peptides, HLA-II and TcRs that should be evaluated to characterize this interaction system are staggering. This calls for high-throughput and bioinformatics-based approaches. Indeed, bioinformatics has been a driving force in describing MHCII specificity. Based on binding and affinity data, several different algorithms have been developed capable of predicting peptide MHC-II interactions (e.g. the NetMHC suite) as an efficient approach to address the vast number of peptides and MHC allotypes. Prediction algorithms powered by artificial neural networks (ANNs) are dependent on large bodies of data which are costly to generate since each peptide conventionally needs to be individually synthesized and handled before conducting binding experiments. More recently, proteomics-driven, high-throughput identification of peptides eluted off MHC-II molecules has become possible and used to improve the performance of MHC-II prediction algorithms (2).

While the prediction tools available today have improved significantly over the years, they are still impaired by a high false positive rate (3) making it difficult to select true T cell epitopes. This is recognized as a major current problem in the development of personalized medicine and immunotherapies, in particular those modes that are based on real-time discovery of tumor neo-antigens (4, 5). The recent advances within the field of cancer immunotherapy have suggested that immunization with tumor derived neo-antigens is a very promising strategy and growing evidence suggests that MHC-II-restricted neo-antigens should be included (4, 6, 7). Hence, there is an increasing demand for more accurate and efficient predictors of peptide MHC-II interactions, underscored by the pharmaceutical industry now focusing on developing MHC-II prediction tools (8).

We have for years chosen a biochemical approach to study peptide MHC interactions (9). This has resulted in the generation of a large collection of recombinant HLA molecules, which have been used to generate tens of thousands of data points subsequently used to develop the NetMHC suite of prediction algorithms (10–12), which continues to be updated. While high-throughput assays have been developed to study peptide-MHC interactions, classical orthogonal peptide synthesis remains a very costly and time-consuming bottleneck. However, within the last decade, high-density peptide microarray platforms, potentially containing up to millions individual peptides/array, have become available at a cost as low as a few cents/peptide. We have recently developed and used high-density peptide microarrays to characterize linear sequence motifs such as enzyme cleavage sites and B cell epitopes in great detail (13–18). Such peptide microarrays containing C-terminally anchored 15-mer peptides should in theory be able to interact with MHC-II molecules due to its open-ended peptide-binding cleft allowing peptides to protrude through the ends of the cleft; something that should allow C-terminal tethering.

The ability to acquire millions of peptide-MHC-II binding data in a single experiment could potentially transform CD4+ T cell epitope discovery. In particular, the combination of such data with machine-learning methods has the potential to transform the development of peptide-MHC-II predictors. Here, we have investigated the binding of HLA-DRA1*01:01/HLA-DRB1*01:01 and HLA-DRA1*01:01/HLA-DRB1*03:01 to high-density peptide microarrays containing ∼70k random peptides in triplicate; used the binding data to train ANNs, validated these using peptides eluted off HLA-DRA1*01:01/HLA-DRB1*01:01 and HLA-DRA1*01:01/HLA-DRB1*03:01, and compared the prediction power to ANNs trained on public available binding affinity data from the IEDB (19).

## Results

We investigated the binding of HLA-DR molecules to peptides, synthesized *in situ* on microscope glass slides (17), by *de novo* folding of recombinant HLA-DR molecules (20) followed by staining with a monoclonal mouse anti-HLA-DR antibody known to react with a conformational HLA-DR alpha epitope, exclusively expressed by correctly folded HLA-DR alpha-beta hetero-dimers (21) and finally stained with Cy3-labeled polyclonal goat anti-mouse IgG. Only in the presence of both HLA-DRA and HLA-DRB did we obtain a staining pattern corresponding to the location, geometry and size of relevant peptides synthesized *in situ* (figure 1) confirming that conformationally intact peptide-HLA-DR complexes were obtained. We found no binding of the Cy3-goat anti-mouse IgG to the peptide arrays in absence of HLA-DRA, HLA-DRB, or L243 (figure 1) showing that the signals obtained was dependent on the presence of *on chip* generated peptide-HLA-DR complexes.

**Figure 1.**
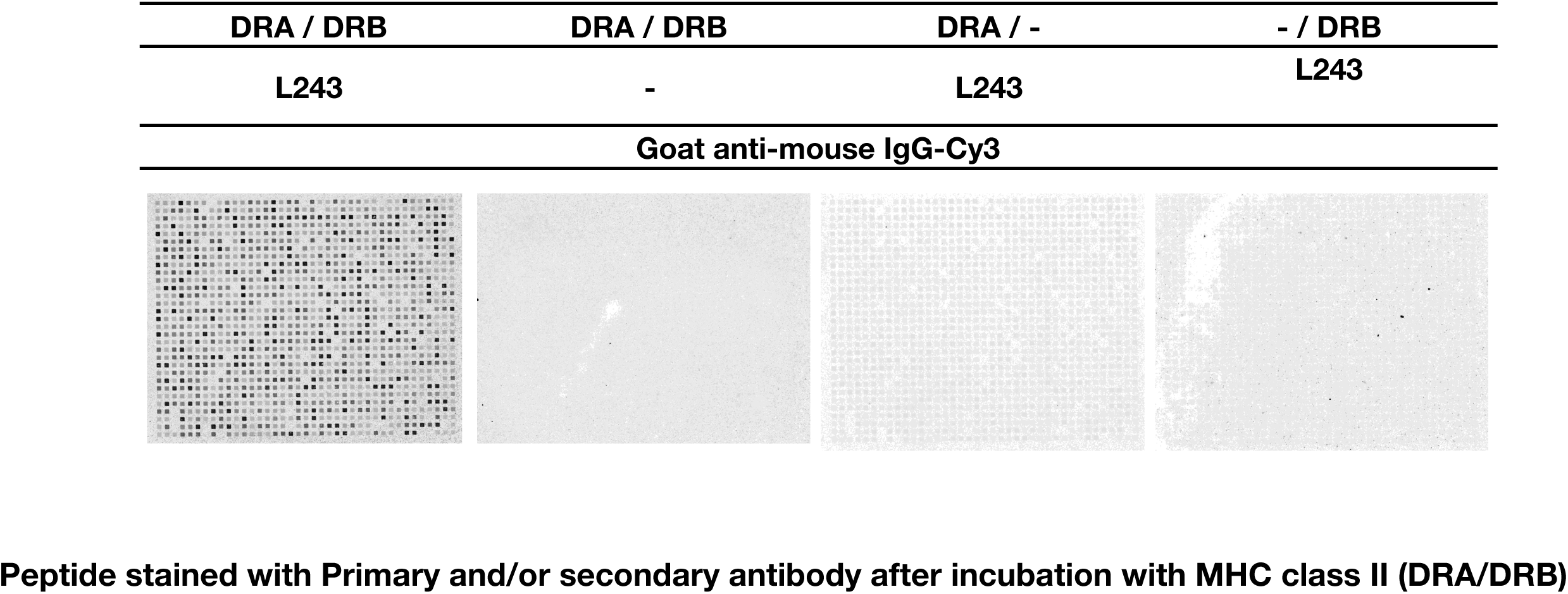
Peptide microarray incubated with HLA-DRA1*01:01 / HLA-DRB1*03:01 stained with monoclonal anti-HLA-DR (L243) and Cy3-conjugated goat anti-mouse in combinations indicated in the figure heading. A fluorescent signal was only detectable in presence of HLA-DRA, HLA-DRB, mouse anti HLA-DR and Cy3-goat anti-mouse suggesting that the binding assay reveals HLA-DR molecules bound to specific peptides.

We determined the optimal peptide length to be synthesized on these peptide microarrays by synthesizing peptides with lengths from 9-15 amino acid residues and using the binding data to train artificial neural networks. Initially, NNAlign-1.4, was used to evaluate the effect of peptide length on network performance expressed as PCC and RMSE. with an optimal performance identified as maximizing the PCC in combination with the lowest RMSE. This identified a peptide length of 13 as being optimal for developing efficient predictors with a combination of high PPC and low RMSE. It should be noted that later versions of the algorithm such as NNAlign-2.1, which allows the incorporation of insertions and deletions in the analysis, can handle longer length (at least up to 15 aa) without loss of performance (suppl. figure 1).

A list containing 63,802 peptides was generated from random sampling 13-mer peptides originating from an in-house database of human pathogens and synthesized in triplicate on a microscope slide with a randomly distributed localization. The microscope slide was incubated with HLA-DRA1*01:01 / HLA-DRB1*03:01 and stained with L243 / Cy3-goat anti-mouse IgG before obtaining a laser scanning image at 1µm resolution (figure 2). The individual peptide fields (20×20 µm) are clearly distinct in the zoomed section (figure 2, insert) and present themselves with varying intensities reflecting the amounts of peptide-HLA-DR complexes present on each field.

**Figure 2.**
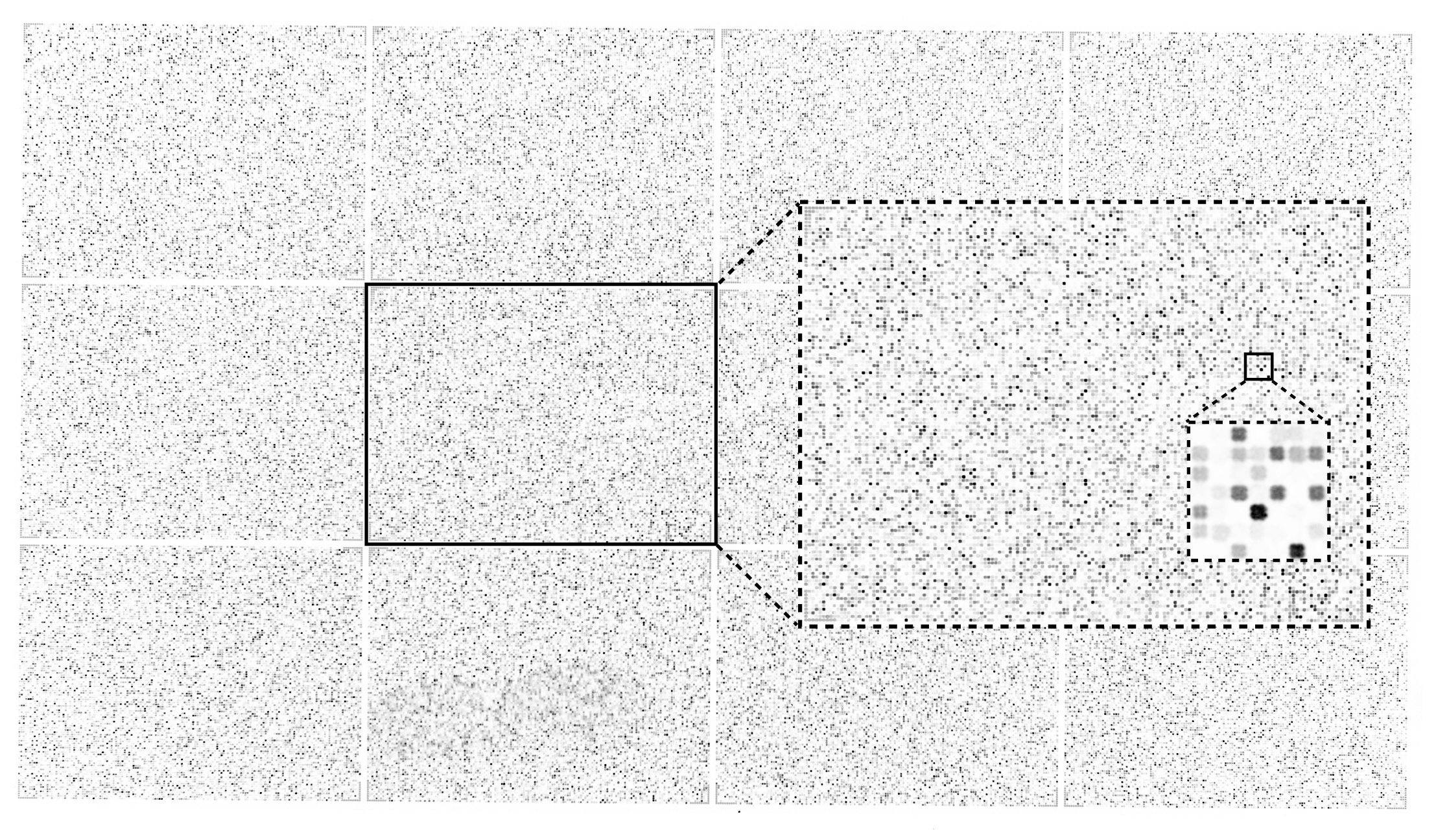
A full view of HD peptide array with 217,000 individual peptides (Area 20×11 mm) divided in 12 virtual sectors. Zoomed inserts of sector 6 and a further zoom of a subsection of sector 6 with individual peptide fields visible. The size of each peptide field is 20×20 µm with a 10 µm pitch scanned with a resolution of 1µm/pixel.

Each individual peptide field was quantified with a proprietary software (PepArray, Schafer-N, Denmark), and the mean signal and the coefficient of variance (cv) based on triplicate values was calculated for each peptide. For both HLA-DRB1*01:01 and HLA-DRB1*03:01, we found a high concordance between triplicate signal values generally showing cv<0.25 for peptides with mean signal values > 50% of the max signal. Peptides with mean signal in the lower 10% of max value showed a marked increase in cv suggesting that these lower signals approached the noise level of detection (figure 3A). Inclusion criteria for downstream ANN training was chosen at cv < 0.5, which almost exclusively filtered away weak and non-binding signals. The mean signal of the remaining (cv<0.5) peptides were, for both HLA-DR allotypes, displaying a log-normal distribution (figure 3B); for ANN training signal values were log-transformed to bring the signal in the range [0-1] and reduce the distribution skewness, figure 3C.

**Figure 3.**
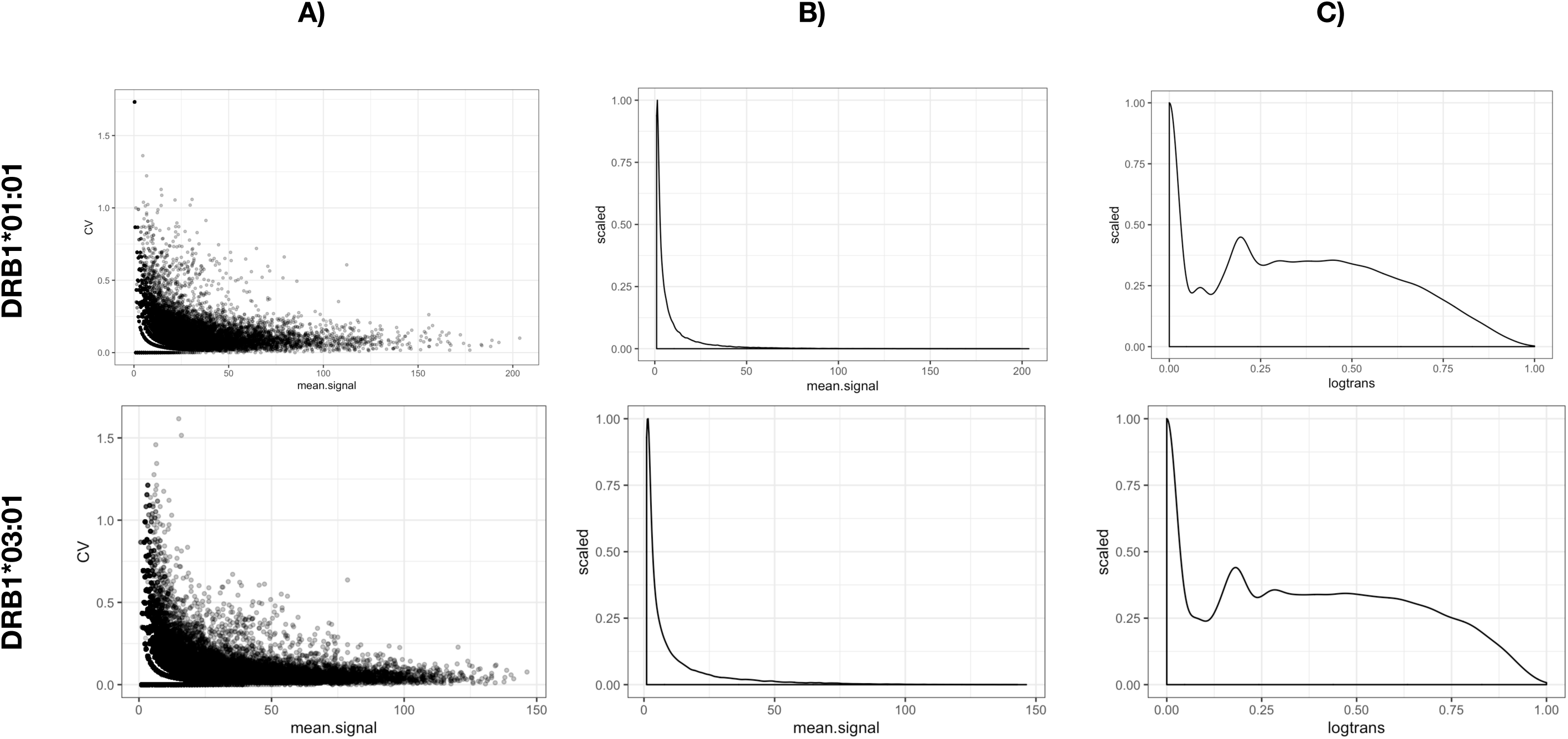
A) Raw data, Mean Signal versus coefficient of variation (cv) calculated from triplicate signal values. B) Signal distribution of the calculated Mean Signal. C) Signal distribution after Log transformation of the mean signal calculated after optimization with a BoxCox transformation (R package EnvStats Version 2.3.1).

By comparing the log-transformed signals of HLA-DRB1*01:01 against HLA-DRB1*03:01 peptide by peptide we found, as expected, the two molecules to bind to very different peptides (figure 4, Left), confirming that a particular HLA-DR molecule has preference towards different subset of peptides. This is further confirmed when considering only the subset of data with log-transformed signal > 0.5 (figure 4, left red square) for both molecules. Here the Spearman rank correlation was SRC = 0.11 suggesting a very limited specificity overlap between the two molecules for strong binding peptides. The reproducibility of the HLA-II/peptide microarray binding assay was examined by repeating the peptide array synthesis (i.e. the peptide design and layout was identical to the first peptide microarray) and repeating the binding of DRA/HLA-DRB1*01:01 and DRA/HLA-DRB1*03:01. Plotting the log-transformed signals from duplicate binding experiments against each other by HLA, we found a Pearson correlation coefficient (PCC = 0.92) for both HLA-DRB1*01:01, HLA-DRB1*03:01(figure 4, middle and right) suggesting that the binding of each HLA-DR molecule was highly reproducible. To assess whether the binding data possessed peptide sequence-based information of a quality sufficient to support the development of ANN predictors, we submitted the data-sets from HLA-DRB1*01:01 and HLA-DRB1*03:01 to the publicly available NNAlign server with the network architecture parameters specified in table 1. As a reference and comparison to the peptide microarray data, we used the original data sets (originating from IEDB binding data) used to train the NetMHCII prediction algorithms (available at https://services.healthtech.dtu.dk/service.php?NetMHCII-2.3) and subjected them to NNAlign with the exact same network architecture as the peptide array data sets (table 1). For both HLA-DRB1*01:01 and HLA-DRB1*03:01, ANN training with the relevant peptide microarray data-sets returned prediction models with a very high internal correlation (PCC > 0.95) between predicted and observed data (figure 5). The results, summarized in table 2, suggests that the peptide microarray data-set of the respective HLA-DR molecules contains a distinct and recognizable pattern that can be extracted by the NNAlign algorithm.

**Figure 4.**
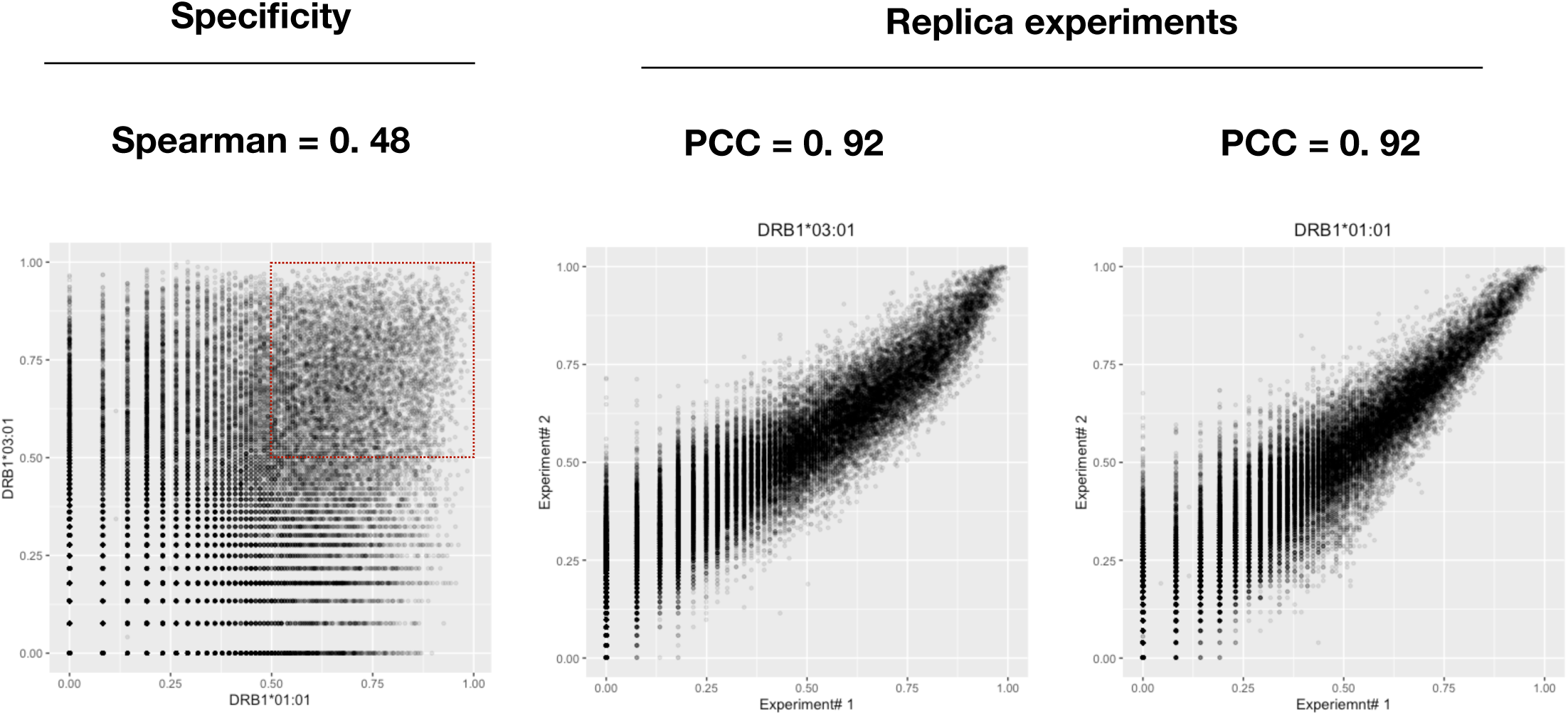
Left: Log transformed signals of DRB1*01:01 plotted against DRB1*03:01 by peptide sequence show a weak correlation (PCC = 0.47). Considering peptides with a predicted score>0.5 returns a SRC=0.11, suggesting HLA-DR specificity. Middle and Right: Log transformed signals obtained by replica experiments of DRB1*03:01 and DRB1*01:01, respectively, correlates well (PCC = 0.92) conferring an acceptable reproducibility.

**Figure 5.**
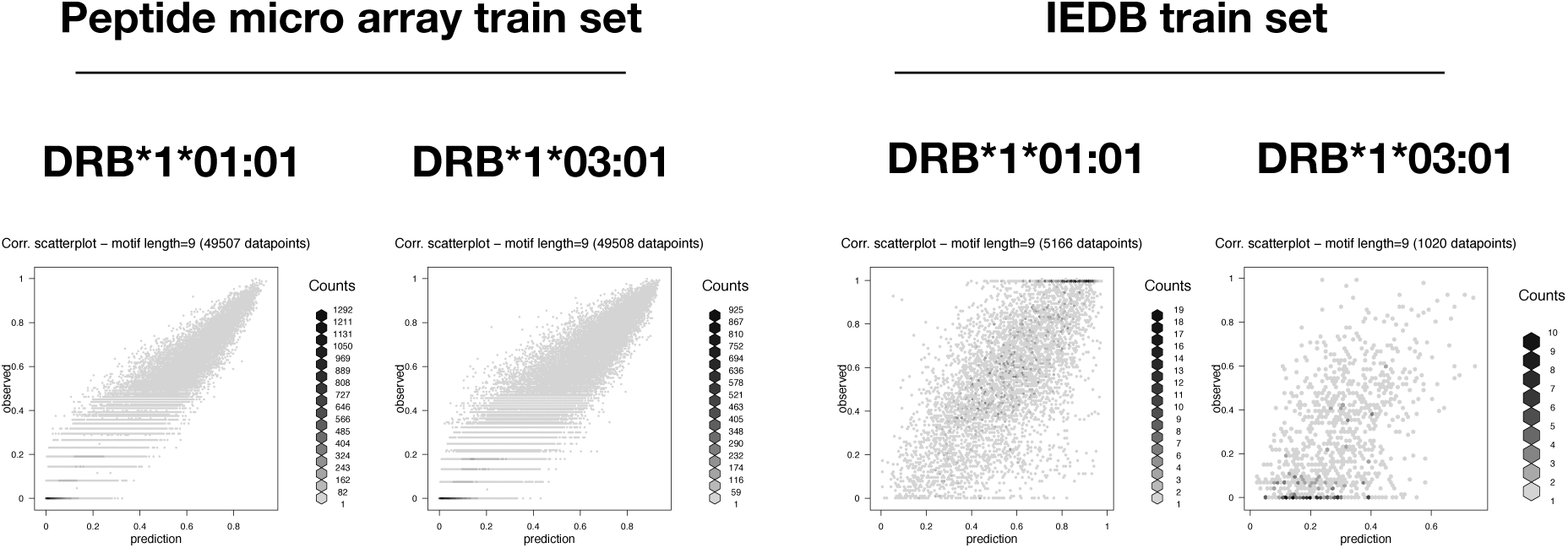
Scatter plots of predicted versus observe values (log transformed signals) of NNAlign-2.1 models trained on ∼50k peptide sequences obtained from peptide microarrays with SPARSE encoding. IEDB train sets (downloaded as pairs of peptides sequences and log transformed binding data from https://services.healthtech.dtu.dk/service.php?NetMHCII-2.3) encoded with SPARSE. The internal model performances are listed in table 2. BLOSUM encoding rendered network ensembles with comparable performances (data not shown). The neural network architecture is disseminated in table 1.

**Table 1.**
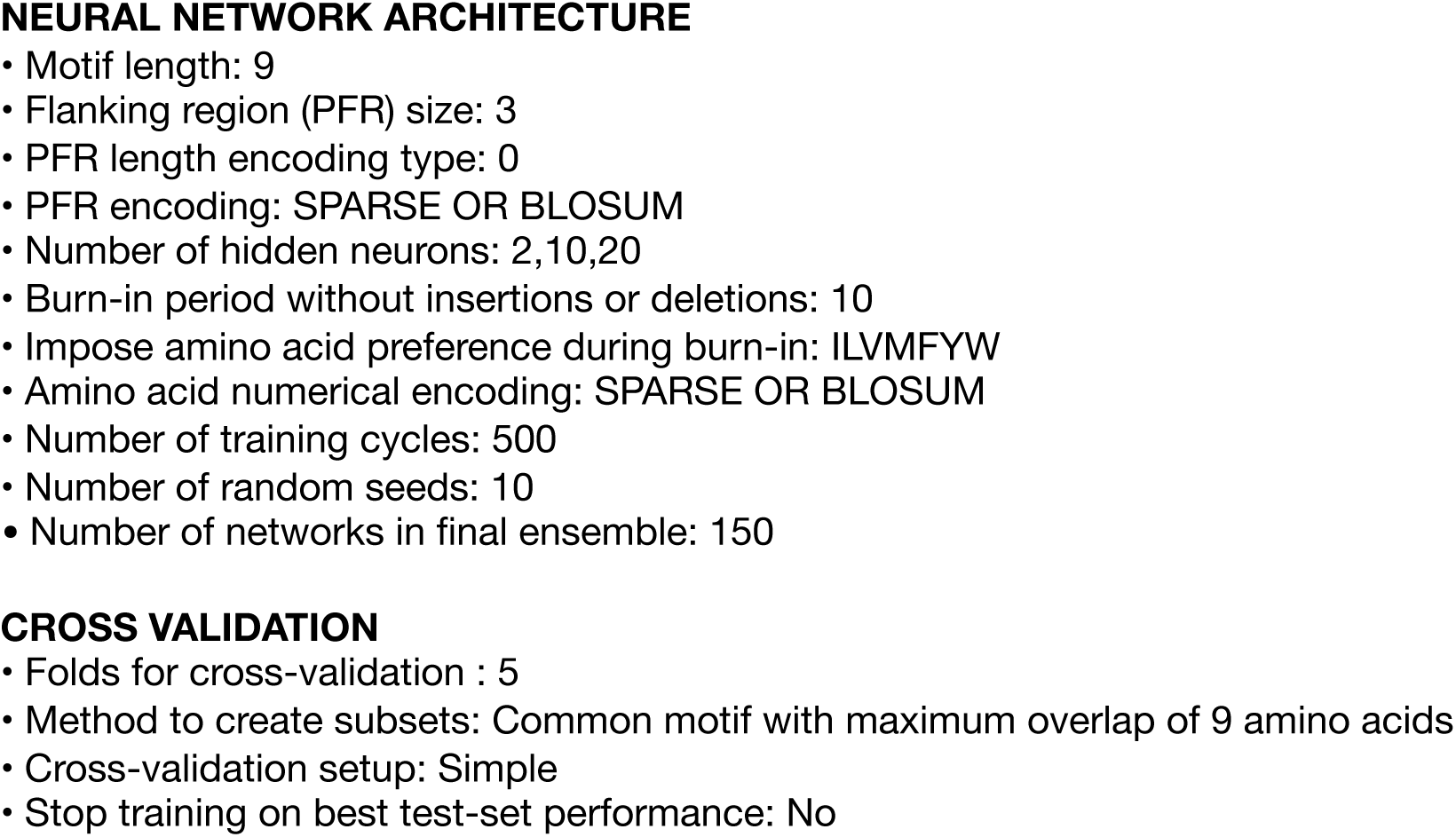
ANN summary. Summary of NNAlign architecture and encoding applied to all four models. Data sets were reduced to 50,000 peptide sequences according to the maximum accepted number of inputs by the server.

**Table 2.**
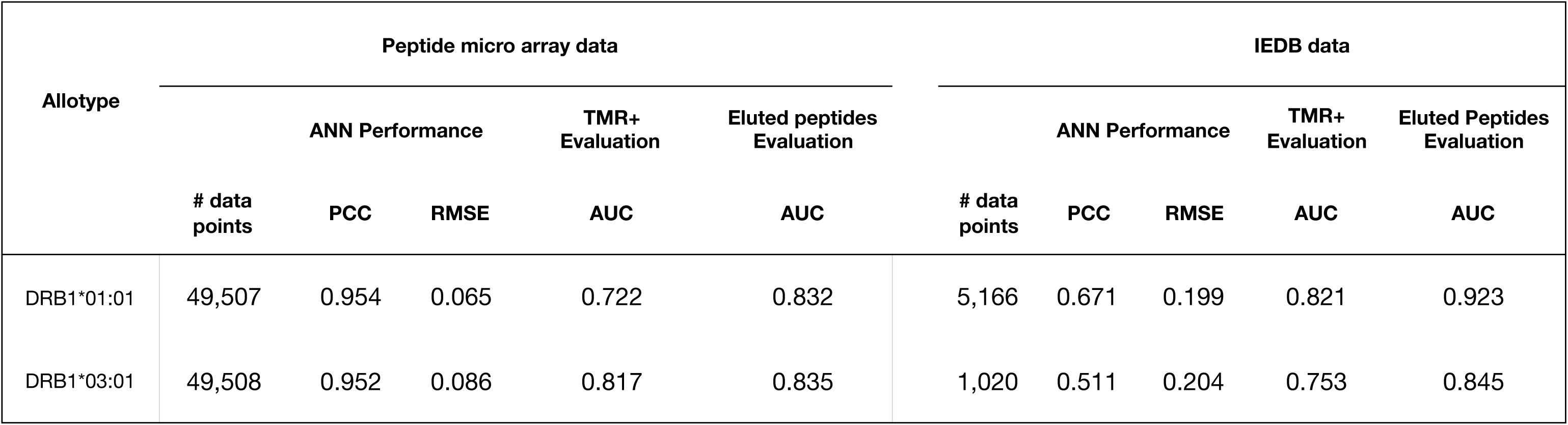
ANN output and Performance. Summary of the final network ensemble for all models listing the number of data points, prediction performance reported as Pearson correlation coefficient (PCC) and Root mean square error (RMSE). The mean AUC values are Area under the curve reported by a ROC analysis of predicted scores of IEDB epitopes versus epitope source antigens.

The sequence logo representations of the final NNAlign network ensembles trained on peptide microarray data sets revealed motifs with distinct anchor residues in positions P1, P4, P6 and P9 for both HLA-DRB1*01:01 and HLA-DRB1*03:01 (figure 6). Although P1 in HLA-DRB1*01:01 did not appear to be as prominent an anchor position (reflected by a relatively low bit score) as normally observed for P1 in many HLA-II motifs, we found the motifs and anchor positions to be comparable to the commonly accepted motifs of the respective molecules. A head to head comparison of the final DRB1*01:01 ANN network ensembles trained on peptide microarray versus network ensembles trained on IEDB data, showed overall that the peptide microarray generated motif appeared more blurred with less distinct anchor positions (P1 in particular). In contrast, the motif for HLA-DRB1*03:01 as derived from the microarray data appeared sharper and with higher information context compared to the motif derived from the IEDB data.

**Figure 6.**
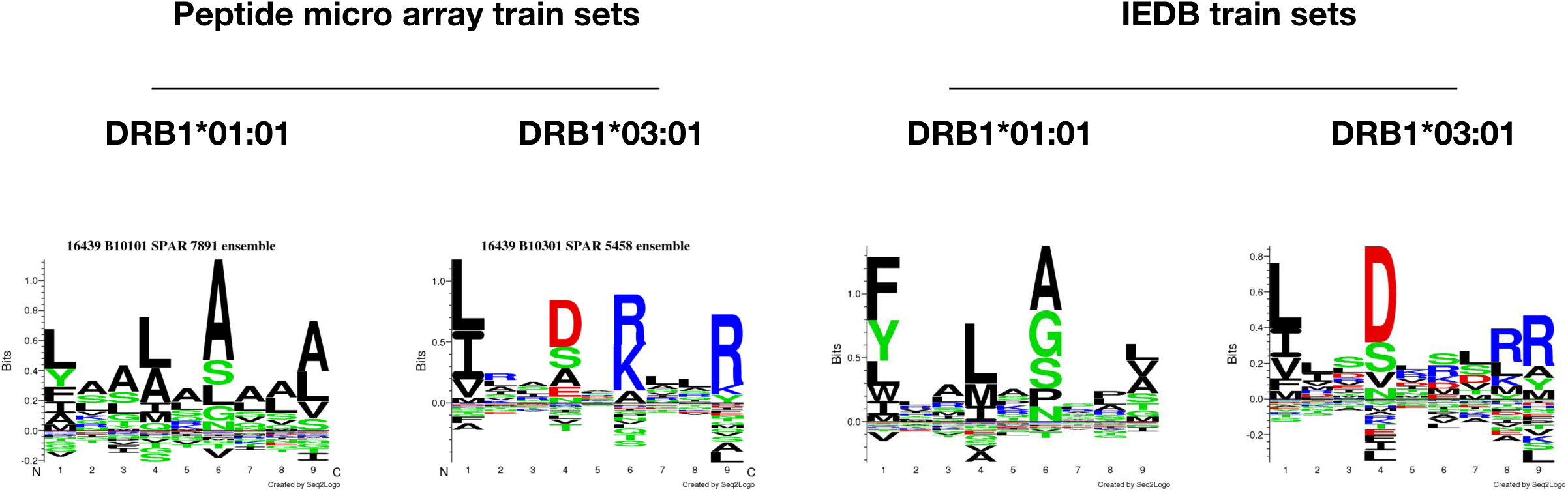
LOGO representation of the final NNAlign network ensemble based one the peptide microarray data set and IEDB data sets trained with the neural network architecture listed in table 1.

To benchmark the prediction power of the final network ensembles, we evaluated the prediction of HLA-DRB1*01:01 and HLA-DRB1*03:01 epitopes extracted from IEDB with selection criteria outlined in table 3 (note that epitopes were curated for cases where the epitope sequence not being present in the corresponding source antigen). To benchmark the peptide microarray driven models which operates in a 13-mer peptide space, each source antigen was *in silico* digested into overlapping peptides with the length of the epitope; within each of these peptides, the highest predicted score of the underlying overlapping 13-mers was assigned to the peptide. Similarly, for each epitope was the highest predicted 13-mer within each epitope assigned to the epitope. The IEDB data set driven models were trained on peptides with varying lengths, hence predictions were made on the epitope sequence, and overlapping peptides from the source antigen with same lengths of the epitope. By comparing the prediction of each epitope against the predictions of the overlapping source antigen peptides, an AUC was calculated for each epitope; AUC values of each epitope and models are shown in figure 7. The overall performance (mean AUC values) of each model, summarized in table 2, shows that the prediction power of IEDB data driven networks is higher (p<0.05), AUC=0.821 as compared to AUC=0.722 of peptide microarray driven networks, with respect to predicting HLA-DRB1*01:01 epitopes. In contrast. was the power to predict HLA-DRB1*03:01 epitopes higher (p<0.05) by the peptide microarray driven networks AUC = 0.817 compared to IEDB driven networks predicting with an AUC = 0.753. As a further validation, we evaluated the prediction of peptides eluted off HLA-DRB1*01:01 or HLA-DRB1*03:01, and sequenced by mass spectrometry (22) and as described here (eluted peptide sequences provided in Suppl. Table 1). Peptides (5515) of different lengths eluted off HLA-DRB1*01:01 were in silico digested to length 13 amino acids and subjected to prediction, the highest predicted 13-mer within each eluted peptide were used in the evaluation against 5000 randomly generated 13-mer peptides. An identical strategy was applied for the (5713) eluted HLA-DRB1*03:01 peptides evaluated against the same 5000 randomly generated peptides. Density plots (figure 8) of the predicted scores confirmed that all four network ensembles are assigning higher scores to peptides eluted off relevant MHC-II allotypes than to random peptides, resulting in a clear skewness of the score distribution. The AUC values, summarized in table 2, of the evaluation of eluted peptides against random peptides were comparable to AUC values obtained for the evaluation of epitopes. Using a student’s t-test to compare the performances of the peptide microarray driven models against the IEDB driven models, we found no statistical difference (p=0.17, p=0.31, HLA-DRB1*01:01 and HLA-B1*03:01, respectively) of the combined mean AUC of epitopes and eluted peptides. A direct comparison of the prediction scores of eluted peptides between the peptide microarray driven models and IEDB models (fig 9) reveals that, the peptide microarray and IEDB model performances are comparable, with Spearman rank correlations (HLA-DRB1*01:01) SRC=0.66 and (HLA-DRB1*03:01) SRC=0.71. There is a tendency towards agreement between the models in particular for peptides with high prediction scores. It is possible that the nature of the binding data, which is obtained from two different assays, chip binding and traditional affinity data (IEDB), contributes to the differences. For all data sets, the peptide with the strongest measured binding is transformed to a value of 1 in the training sets. Hence, the differences in distribution of the transformed data originating from the different data sets may in part explain some of the differences observed for prediction scores < 0.8; which in any case translates into weak to non-binders and thus less relevant.

**Figure 7.**
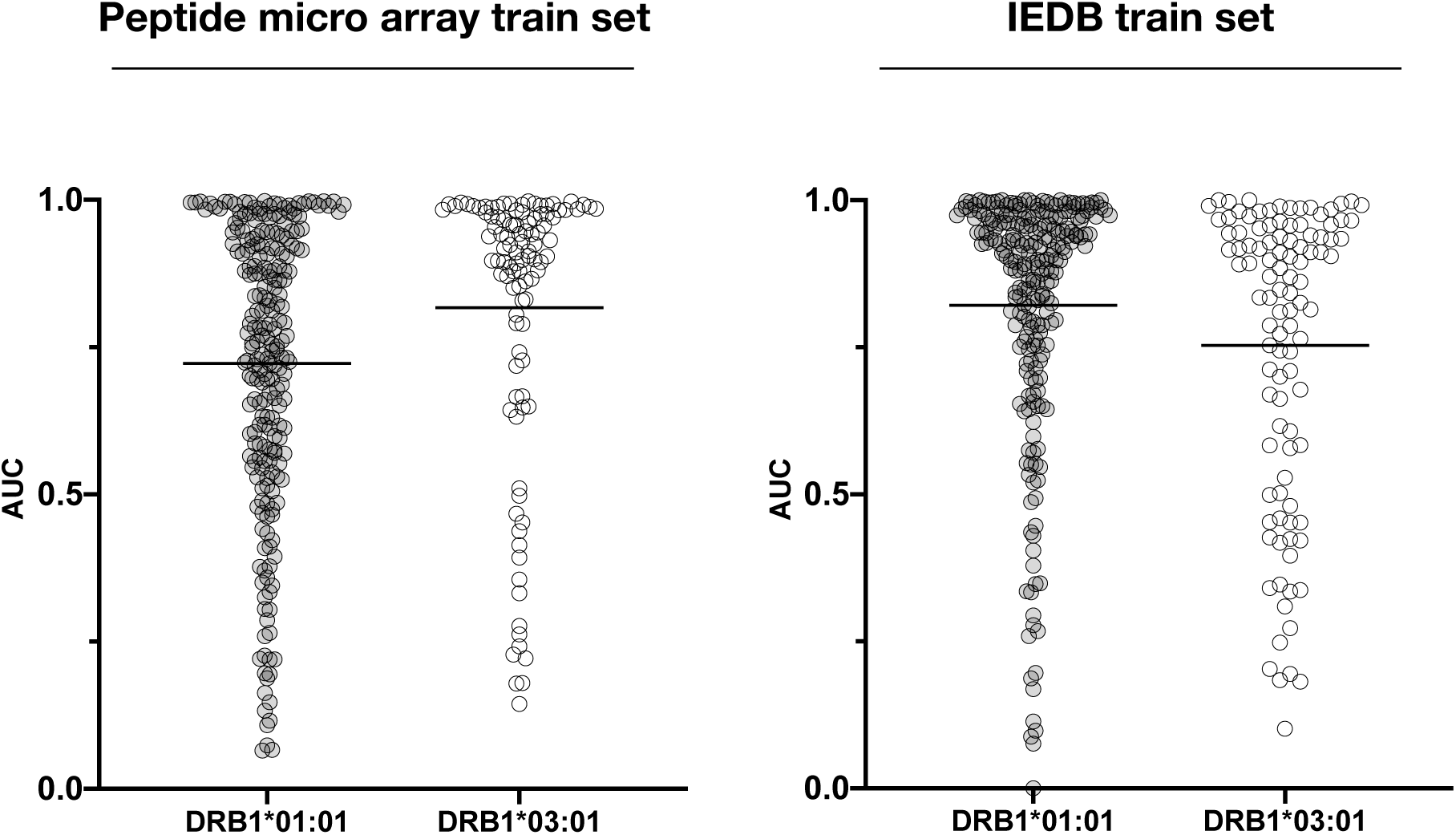
AUC values of HLA-DRB1*01:01 (light grey) and HLA-DRB1*03:01 (white) epitopes based on predictions by network ensembles trained on peptide micro array data (left) and IEDB binding data (right). Neural network architecture is listed in table 1 and the mean AUC values are listed in table 2. In general, all four models assign high prediction scores to peptide sequences which can be identified as HLA-II tetramer validated epitopes as expected if the input data reflects the peptide HLA-DR interaction. The prediction performances reported as AUC values are comparable between the models with respect to allotype. This indicates that the data obtained by peptide microarray binding studies contain specific peptide-HLA-DR information comparable to traditional binding data and suitable to ANN training.

**Figure 8.**
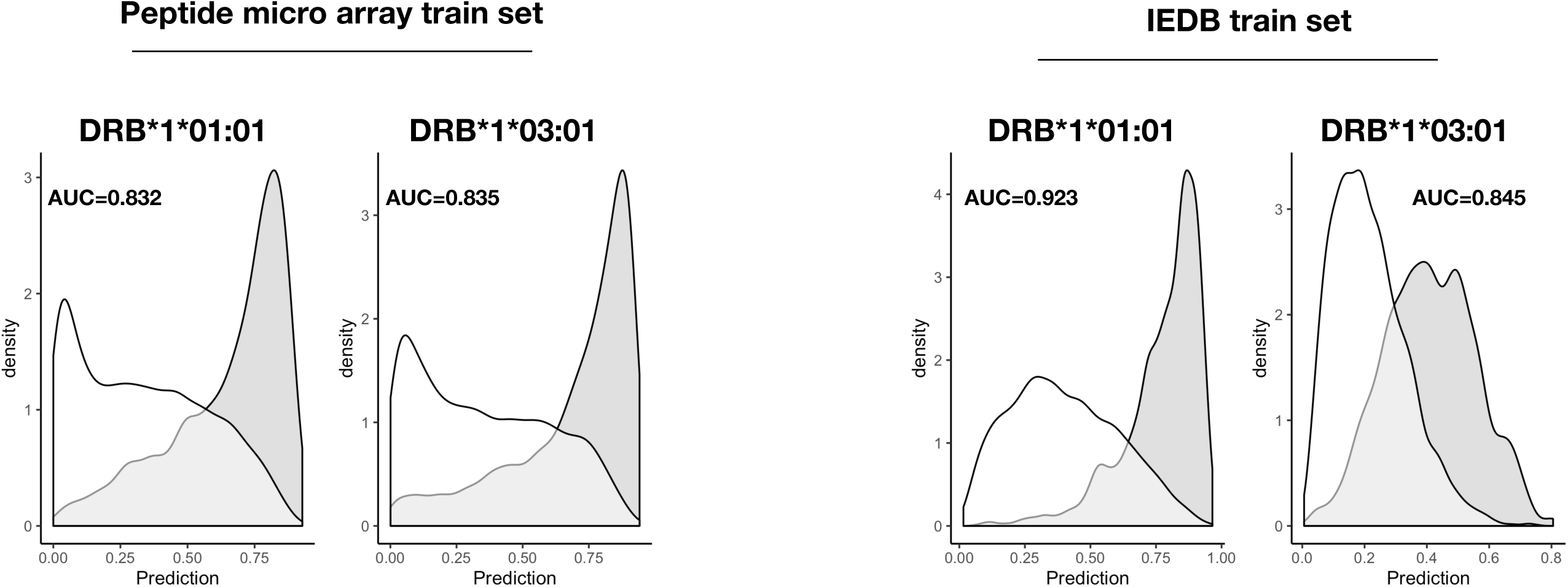
Predicted scores of peptides eluted off HLA-DRB1*01:01 and HLA-DRB1*03:01 and compared to random peptides. Left, the peptide microarray model prediction of eluted peptides (grey density) are clearly skewed towards high scores while random peptides (white density) primarily are skewed towards low scores for both HLA-DR allotypes. Right, predicted scores by the models trained in IEDB data presents a similar pattern of eluted peptides (grey density) and random peptides (white density). The model performances are summarized in table 2 showing that HLA-DRB1*03:01 models performed at par while the HLA-DRB1*01:01 model based on IEDB data sets had a better prediction power compared to the model trained on peptide microarray data.

**Figure 9.**
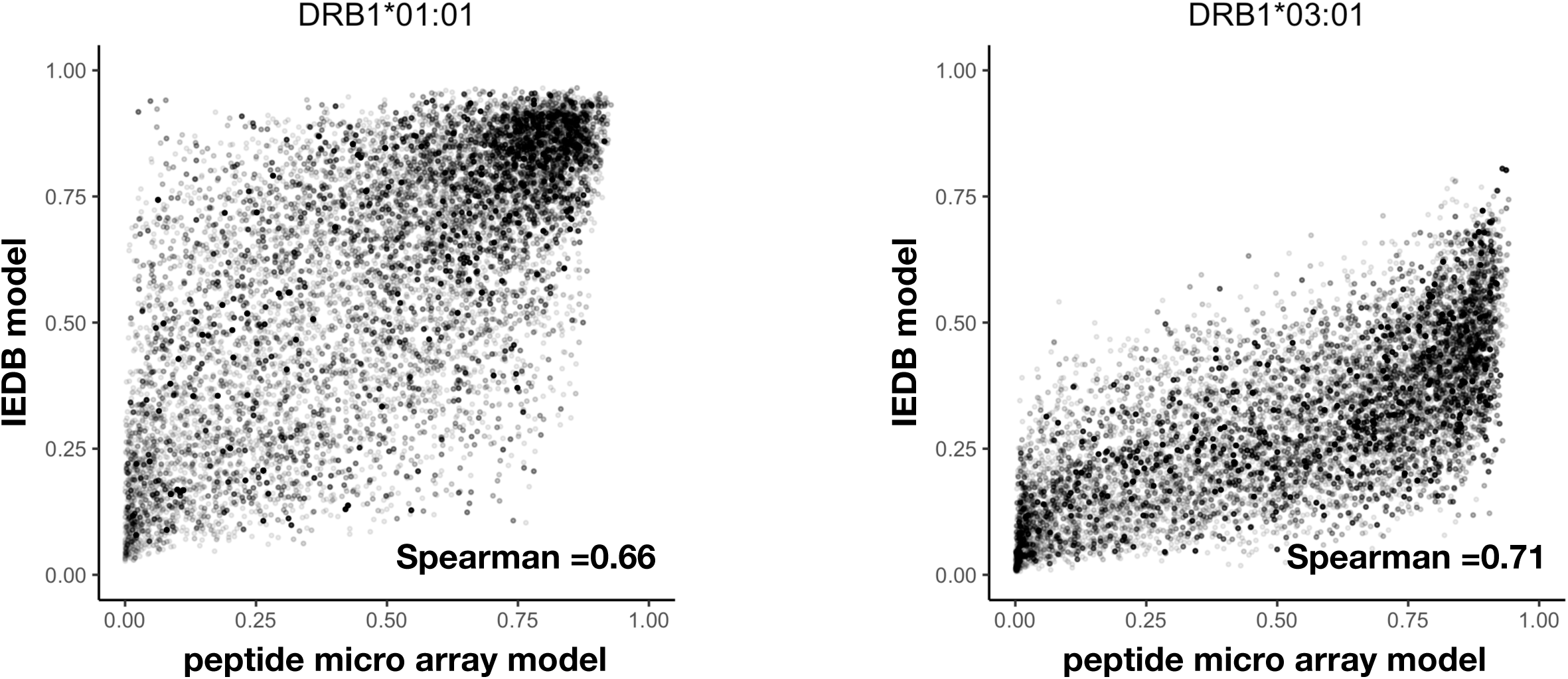
Scatter plots of eluted peptide scores (Left: HLA-DRB1*01:01, Right: HLA-DRB1*03:01) predicted by peptide microarray models versus IEDB models. The Spearman rank correlation of the respective data-set are inserted.

**Table 3.** Evaluation data. Evaluation data sets (epitopes) extracted from IEDB after selecting positive assays (tetramer /multimer staining).

| Source | HLA | Applied filters | # of Epitopes | # of Epitopes curated | # of Source Antigens |
| --- | --- | --- | --- | --- | --- |
| IEDB | DRB1*01:01 | • Positive Assays Only<br>• Qualitative binding (MMR/TMR)<br>• No B cell Assays<br>• No MHC ligand assays<br>• MHC restriction - ID MRO_0001279 HLA-DRB1*01:01 | 266 | 235 | 80 |
| IEDB | DRB1*03:01 | • Positive Assays Only<br>• Qualitative binding (MMR/TMR)<br>• No B cell Assays<br>• No MHC ligand assays<br>• MHC restriction - ID MRO_0001284 HLA-DRB1*03:01 | 114 | 105 | 49 |

Collectively, the high internal performance of the peptide microarray driven models and a prediction power comparable to prediction models based on traditional binding data (IEDB), suggests that high-density peptide microarrays can be used to generate relevant peptide-HLA-II binding data. As a final examination of the prediction models, we calculated the distance between the models one-by-one and their individual distances to NetMHC-2.3, summarized in table 4. Briefly, the distance is calculated as 1-SRC^2^, where SRC is the Spearman rank coefficient found by comparing prediction scores of 100,000 random peptides from two prediction models. We found the distances of network ensembles to be closer within allotypes (HLA-DRB1*01:01 distance = 0.31 and HLA-DRB12*03:01 distance = 0.43) than between allotypes (0.49 < distance < 0.85) underscoring that the training data contains allotype specific binding data.

**Table 4.**
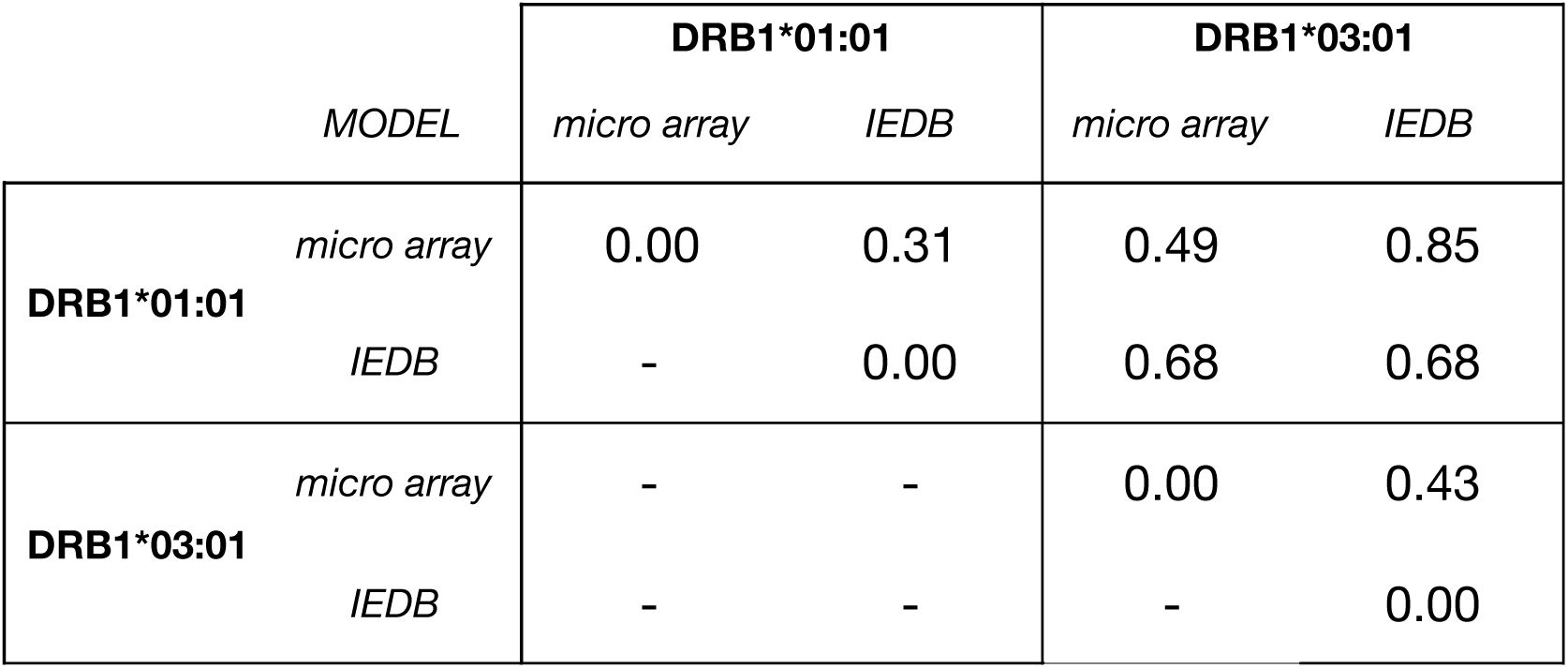
Distances between predictions models. Table of calculated distances between the models trained on peptide microarray data, IEDB binding data and NetMHC-2.3. The distance is calculated as 1-R^2^, where R is the Spearman rank correlation calculated on the prediction scores of 100,000 random peptides.

In conclusion, the binding data obtained from HLA-II-peptide microarray assays produced comparable HLA-II peptide prediction models compared to models trained on peptide-HLA-II affinity data available at IEDB, and as such represents an attractive way forward to generate large amounts of peptide-MHC-II binding data and to improve existing prediction models.

## Discussion

CD4+ T cells arguably perform the most important function of the immune system; by controlling and coordinating the responses of most immune cells they essentially orchestrate the overall specificity and reactivity of the immune system. To this end, they survey peptide-MHC class II complexes and determine whether these peptides are of self or non-self-origin. This essential process of peptide sampling and display is diversified immensely by the polymorphism of the MHC system, which assures that no two individuals will perform self-non-self-discrimination in an identical manner thereby avoiding the evolution of pathogen variants that can escape the immune systems of the entire species. An in-depth understanding of which peptides bind to MHC-II and how this interaction is affected by MHC polymorphism is of paramount importance for our ability to understand and manipulate the immune system. Many autoimmune diseases (e.g. type I diabetes, multiple sclerosis, rheumatoid arthritis, narcolepsy and many more) are tightly associated with specific MHC-II allotypes, which obviously make these allotypes interesting in their own right. More recently, the field of personalized immunotherapy - emerging as promising cancer treatments – increasingly rely on a better understanding and prediction of immune specificities in a much broader representative coverage of the “MHC space” underscoring the relevance of pan-specific predictors.

A need for experimental coverage became apparent when pan-specific predictors were applied to MHC allotypes lacking experimental data and it could be shown that these pan-predictors are more accurate when they have been trained on experimental data from a closely related allotype rather than a more distant allotype (23). The development of these bioinformatics methods requires large-bodies of data that are representative of peptide-MHC-II interactions. In other words, it requires access to large panels of natural and/or synthetic peptides representing peptide-MHC-II binding events. By combining recombinant MHC-II molecules with high-density peptide array technology, it is be possible to generate the large bodies of binding data representing many different allotypes, and do so at significant cost reductions compared to generating the same amount of data using more traditional binding experiments.

Here, we have demonstrated that MHC-II can interact with *in situ* synthesized peptides in a high-density array format; that the peptide MHC-II interaction is specific; that there is a systematic binding pattern which is interpretable by ANNs, here exemplified by NNAlign training (24, 25); furthermore the resulting prediction models are able to predict peptide MHC-II binding at a level comparable to that of models trained on traditional binding data deposited at the IEDB which have been used to develop NetMHCII models (25). We also found the binding motifs revealed by the models trained on peptide microarray data comparable with the motifs obtained by training on IEDB binding data.

The capacity of this high-density peptide microarray technology should enable large-scale analyses of any MHC-II allotype of interest even including all post-translational modifications that are available for solid-phase peptide synthesis (SPPS). Without the limitations inherent to conventional SPPS, peptides could be selected entirely based on the scientific and/or practical merits of the question at hand. A completely random selection of peptides, as applied here, would ensure a truly unbiased approach to define MHC-II specificity. Alternatively, one could randomly extract peptides from the proteomes of organism(s) of interest (microbiomes for infectious diseases, the human proteome for autoimmune disease, onco-proteomes for cancer etc.). Other “peptide-intensive” strategies could also be pursued e.g. using an initial random screen to generate a primary predictor and use that to enrich for binders in a secondary experiment. In contrast, the current cost of obtaining peptides for such mapping projects tend to be prohibitive and may lead to the use of existing prediction tools to downsize sampling with the inherent risk of introducing bias leading to an incomplete representation of the relevant sequence space.

The prediction models developed using peptide microarray binding data are at performance levels comparable to prediction models trained on traditional binding (affinity) data. Thus, peptide microarrays could be used to develop large bodies of peptide-MHC-II binding data and/or to complement other methods capable of generating large amounts of binding data (e.g peptide elution and mass spectrometry) (2, 23). This could even include non-standard amino acids representing post-translational modifications that otherwise are scarcer and/or more difficult to obtain. The massive amounts of representative peptide-MHC-II data that can be generated should allow an exhaustive scanning of both the peptide and MHC spaces. This should support the generation of improved pan-MHC-II predictors, which should allow peptide-MHC-II interactions to be evaluated *in silico* cheaper, faster and at an even higher capacity than any experimental approach, even one enabled by the high-density peptide microarray technology, could do. One area may always benefit from an experimental approach: the binding of epitopes derived from cancer neoepitopes identified by genomics or proteomics sequencing of tumor cells vs normal cells. Here, all mutations and corresponding wildtype sequences could potentially be synthesized in multiple length variants and tested experimentally against all the MHC-II allotypes of a given patient.

One current disadvantage of the present peptide microarray technology is the inability – at least with current technologies – to validate the identity and quality of the peptides. Thus, some of the experimental peptide-MHC-II data points might be erroneous. We surmise that as long as such errors are the exception, rather than the rule, then the predictors should ignore these experimental errors. Notwithstanding, it would be a major advance if the quality of the individual peptides on a microarray could be validated through independent physical means e.g. by mass spectrometry.

Other recent technologies have emerged as possible ways of generating large-scale peptide-MHC-II data. One approach is phage expression of peptide libraries followed by MHC-II selection and sequencing (26). This technology does not offer the same degree of control of the peptides expressed; it requires DNA sequencing to identify the peptides involved rather than the simple “identification by look-up” used by the peptide microarray technology; it does not readily allow the incorporation of post-translational modifications; nor does it allow the identification of non-binders which are important for ANN training. Another recent approach, “immunopeptidomics”, involves sequencing of natural peptides eluted of MHC-II by liquid chromatography followed by mass spectrometry (27). Sequencing natural peptides has the huge qualitative advantage that it includes events of antigen processing and includes post-translational modification. Otherwise, it shares some of the same disadvantage of the phage display approach: lack of control of which peptides and post-translational modification are interrogated; and lack of non-binders. Ideally, one should include data obtained from both synthetic and natural peptide, and generate immune-bioinformatics predictors incorporating both kinds of input.

In conclusion, we have demonstrated that high-density peptide microarrays can be used to generate very large numbers of discrete peptide-MHC-II interaction data; something that represents a major advance in the analysis of peptide-MHC-II interactions, and should become instrumental in developing improved bioinformatics methods representing these important interactions.

## Methods

### Production of recombinant HLA-DR molecules HLA-DRA1*01:01, -DRB1*01:01, DRB1*03:01

HLA-DR molecules were produced and purified according to (20). Briefly, this entailed separate *E. coli* expression of each recombinant HLA-DR chain followed by a multiple liquid chromatography purification steps. The purified HLA-DR molecules were stored in 8M Urea, 25 mM TRIS, 15 mM NaCl at −80°C.

### High-density peptide arrays

Nexterion-E microscope slides (Schott, Jena, Germany) were amino-functionalized with a 2% w/v linear copolymer of N,N′-dimethylacrylamide (Sigma-Aldrich) and aminoethyl methacrylate (Sigma-Aldrich) and used as substrate for solid-phase peptide synthesis. High-density peptide arrays were then synthesized on the surface of the amino-functionalized microscope glass slides using a principle of maskless photolithographic synthesis (17, 28) combined with standard Fmoc peptide synthesis modified with the photolabile NPPOC (2-(2-nitrophenyl)propyl oxycarbonyl). A detailed description of the peptide synthesis has been described elsewhere (13).

### HD peptide array design

The array designs were all made with a proprietary software (PepArray, Schafer-N, Denmark) by importing peptide sequences and randomly distributing the sequences across 12 virtual sectors. A design to analyze the effect of peptide length on MHC-II binding ∼2300 15-mer peptides with random sequences were generated *in silico* and subsequently chopped into overlapping 8-14 mers and randomly distributed across the peptide array.

For the generation of peptide MHC-II binding data, 500k 13-mer peptides were generated from a list of pathogen proteins from which 72,000 peptide sequences were randomly sampled and distributed across the peptide array in triplicates.

### Binding of HLA-class II molecules to peptide microarray

The peptide arrays synthesised on microscope glass slides were hydrated in Lockmailer microscope jars containing 15 ml PBS prior to incubation with HLA-DR molecules.

HLA-DR molecules (DRA1*01:01, DRB1*01:01, DRB1*03:01, immunAware, Denmark) dissolved in storage buffer (8M Urea, 25 mM Tris pH 8, 25 mM NaCl) were diluted in 12 ml re-folding buffer (PBS pH 7.4 suppl. with 0.01% Lutrol F68, Glycerol 20% (vol/vol), TPCK, TLCK, PMSF, TCEP) to a final concentration of 500 nM HLA-DRA and HLA-DRB and transferred to the incubation jar containing the hydrated peptide array and allowed to re-fold at 18°C for 48 hours. Following incubation, the incubation jars were drained and the peptide arrays washed 5x in PBS-T (PBS pH 7.4, 1% Tween-20) followed by incubation with 1 µg/ml of monoclonal mouse anti-HLA-DR antibody (clone #L243) in PBS-T for 2 hours. The peptide arrays were washed 5 times in 15 ml PBS-T before a final incubation with 1 µg/ml Cy3-conjugated goat anti-mouse IgG (Bethyl Lab., TX) in PBS-T. A final wash with 5x PBS-T was conducted before transferring the peptide array to PBS for 20-30 min before drying the microscope slide in a spin centrifuge dryer.

### Data acquisition

The peptide arrays were scanned in a microscope slides laser scanner (INNOSCAN 900, Innopsys, France) at 534 nm, resolution 1µm and stored as TIFF-format images. Prior to qualification of the image scans, each image was cropped to include only the peptide array area surrounded by a thin border, subsequently, the cropped images were rescaled to (10000 x 6000 pixels) and saved in PNG-format.

### Quantification of signals (proprietary software, Schafer-N)

The processed images were quantified with a proprietary software (PepArray, Schafer-N, Denmark) by positioning a virtual grid corresponding to the peptide fields on top of the scanned images and quantifying each peptide field. The quantified signals (8-bit) were stored in a text-file format containing corresponding peptide sequence information, peptide field id, quantified signal, row, column and (virtual) sector information. The quantified data were prepared for ANN (NNAlingn) training by calculating the mean value SD and CV (based on triplicate measurements). Inclusion criteria for ANN training set were set at a threshold of cv < 0.5. The NNAlign server (2.1) https://services.healthtech.dtu.dk/service.php?NNAlign-2.1 accepts columns of a text strings and corresponding numerical values [0-1] with a limit of 50,000 inputs, hence the data-sets were reduced to 50,000 by selecting, based on signal strength, the top 2000 peptides and random sampling of the remaining peptides.

### Data transformation

The data were transformed from the 8-bit format [0-255] to [0-1] using a log-transformation optimized by a BoxCox algorithm written in R using the EnvStats package (29)

### ANN Training

Log-transformed data with corresponding peptide sequences (one letter code) were uploaded to the NNAling 2.1 server and trained with the parameters indicated in Table 2.

### ANN model evaluation

Peptide sequences used to evaluate the ANN models were obtained by selecting IEDB epitopes restricted to HLA-DRB1*01:01 or HLA-DRB1*03:01 and selecting only peptide sequences with a positive multimer/tetramer assay (see list of filters applied in Table 3 Evaluation data). The remaining epitopes were further filtered for inconsistencies between the peptides sequence and reported source antigen and subsequent *in silico* digested into 13-per peptide sequences. The source antigen for each epitope were also retrieved based on the Genbank id and *in silico* chopped into peptides with the length of the relevant epitope in question; each of these peptides were further digested into 13-mer peptides where the highest predicted score of each derived overlapping 13-mer were assigned to the epitope or source antigen peptide. The applied filters and number of epitopes and antigens are summarized in Table 3.

### Isolation of HLA DR-bound peptides

Cell pellets from International Histocompatibility Workshop B-LCLs 9022 (COX: DRA*01:02-DRB1*03:01:01) and 9087 (STEINLEN: HLA DRA*01:02-DRB1*03:01:01) (30) were lysed in a mild detergent. Following lysis, peptide-HLA complexes were affinity purified by anti-HLA-DR antibody (LB3.1). Affinity purified HLA molecules and their peptide cargo were separated using reversed-phase chromatography and peptides were subsequently analyzed using mass spectrometry as described (31). In brief, cells were expanded in RPMI-10% FCS and pellets of 109 cells snap frozen in liquid nitrogen. Cells were ground under cryogenic conditions and resuspended in lysis buffer (0.5% IGEPAL, 50 mM Tris pH 8, 150 mM NaCl and protease inhibitors) and cleared lysates passed over a protein A pre-column followed by an affinity column cross-linked with a monoclonal antibody specific for HLA-DR (LB3.1). Peptide-MHC complexes were eluted from the column by acidification with 10% acetic acid. Peptides were isolated using reversed-phase HPLC (Chromolith C18 Speed Rod, Merck) on an Akta Ettan HPLC system (GE HealthCare). Fractions were concentrated and run on an AB SCIEX 5600+ TripleTOF high resolution mass spectrometer. Acquired data was searched against the human proteome (Uniprot/Swissprot v2012_7) using ProteinPilot™ (v5; SCIEX) using following parameters: database: Human proteins from UniProt/SwissProt v2016_12, no cysteine alkylation, no enzyme digestion (considers all peptide bond cleavages), instrument-specific settings for TripleTOF 5600+ (MS tolerance 0.05 Da, MS/MS tolerance 0.1 Da, charge state +2 to +5), biological modification probabilistic features on, thorough ID algorithm, detected protein threshold 0.05. The resulting peptide identities were subject to strict bioinformatic criteria including the use of a decoy database to calculate the false discovery rate (FDR). A 5% FDR cut-off was applied and the filtered dataset was further analyzed manually to exclude redundant peptides and known contaminants.

## Acknowledgements

Funded by European commission (HIPAD project no. 278832), the Independent Research Fund Denmark award DFF – 6110-00644 and the Danish MS Society award A31444)

Mass spec and DRB1*03:01 elution work was supported by grants and a Principal Research Fellowship from the National Health and Medical Research Council of Australia to AWP (NHMRC) (1165490, 1137739). The authors acknowledge the provision of instrumentation, training and technical support by the Monash Biomedical Proteomics Facility. Computational resources were supported by the R@CMon/Monash Node of the NeCTAR Research Cloud, an initiative of the Australian Government’s Super Science Scheme and the Education Investment Fund.

## Conflicts of interests

C. S-N. is affiliated with Schafer-N, provider of HD peptide arrays

T.O. and S.B. (immunAware ApS, commercial provider of HLA monomers and tetramers)

---

**Suppl. Figure 1.**
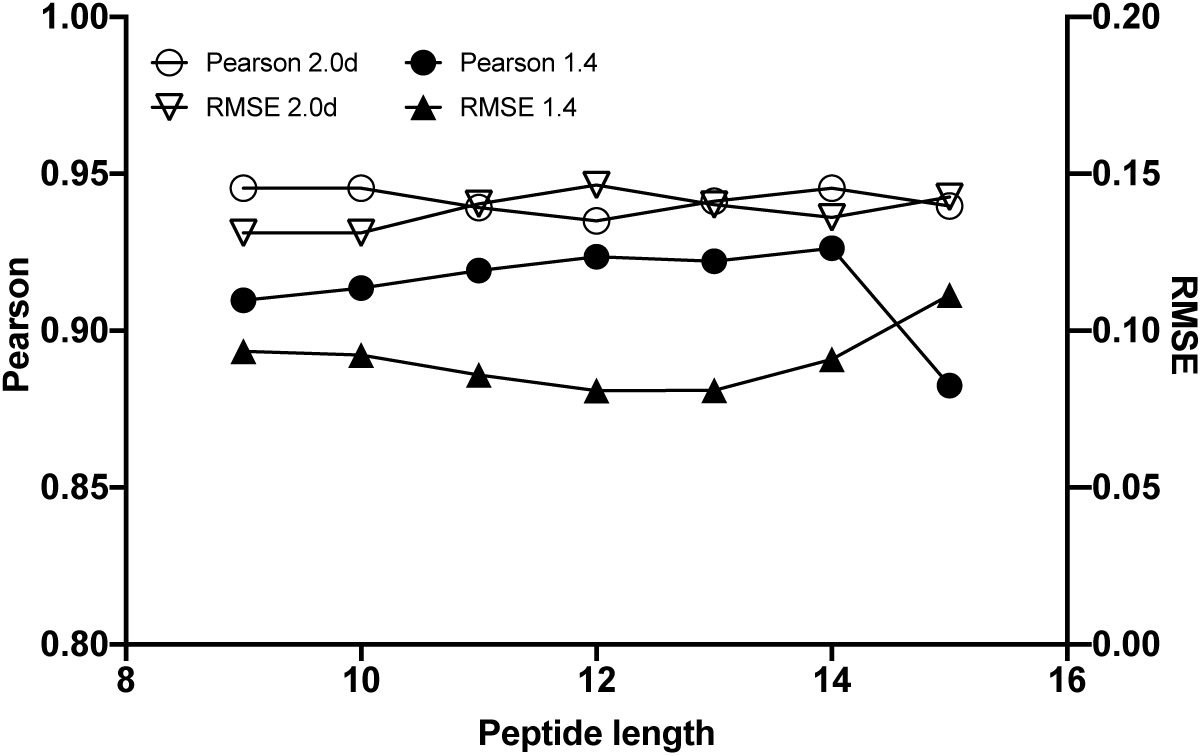
Scanning through different peptide lengths - ANN training (2.0d and 1.4) Optimizing the peptide length. Peptides of lengths 9-15 were synthesised on a peptide array and incubated with HLA-DRB1*03:01. Data from each peptide length were used to train ANNs and the internal correlation and RMSE output from the networks were used to find the optimal peptide length. NNAlign_2.0d (open) shows little or no response to the peptide length input in contrast to NNAlign_1.4 (solid) which clearly show optimal performance at peptide length = 13 (Pearson to RMSE).

